# Radiation-induced persistent DNA damage response and late toxicity in cardiac tissue

**DOI:** 10.1101/2023.02.27.530210

**Authors:** Kay Shigemori, Yanyan Jiang, Jennifer C Martin, Maria A Hawkins, Anderson J Ryan, Eileen E Parkes

## Abstract

**Background:** Radiotherapy treatment is a mainstay of cancer treatments including for thoracic malignancies such as lung or breast cancer. Cardiac toxicity is a recognised long-term complication of thoracic radiotherapy which persists despite improvements in therapeutic modalities. The mechanisms and potential therapeutic targets that could provide cardiac protection in the context of radiation therapy remain incompletely understood. Here we investigated early and late cardiac toxicity following irradiation using the A/J mouse model to identify potential molecular drivers.

**Methods:** Single doses of irradiation of either 13 or 15 Gy were delivered to female A/J mice aged 6 – 8 weeks using a SmART-PLAN system. ECG traces and analysis were performed on anaesthetised mice. Cardiac tissue was harvested up to 32 weeks following irradiation for histological analysis and second-harmonic imaging microscopy.

**Results:** Cardiac RT resulted in cardiac conductivity abnormalities including prolonged QTc interval. Additionally, an increase in pericardial and perivascular fibrosis was noted with a marked increase in pericardial fibrosis at 15 Gy compared to 13 Gy. Persistent DNA damage response was identified in cardiomyocytes at 7 days post irradiation and polarisation of cardiac-infiltrating macrophages towards a CD206+ M2-like phenotype at the same timepoint.

**Conclusions:** Our data introduce the A/J mouse as a novel model for the study of physiologically relevant early and late cardiac toxicities following irradiation. Early events that could contribute to long term toxicities include persistent DNA damage response and repolarisation of macrophages, further investigation of which could identify potential future cardioprotective therapeutic strategies.

## Introduction

Radiation-induced cardiovascular toxicities (RICT) are a potentially serious side effect of radiation therapy (RT). RICT can present clinically in the form of pericarditis, ischaemic heart disease, conduction abnormalities causing arrhythmias, myocardial fibrosis, valvular abnormalities and early mortality^1–3^. The frequencies of radiation-related cardiac events (infarct events and cardiac-related deaths) increase in a dose-dependent manner^4,5^ with no seemingly safe dose of radiation exposure identified. Although technologies to deliver radiation to target tumour tissues while minimising the dose to non-target, healthy tissues have developed in recent years, the risks of RICT persist^6,7^.

Despite the clinical need, few preclinical models are currently available to study radiation-induced cardiac effects^8^. Studying RICT in animal models remains challenging, in part due to the biological differences in the responses to radiation between humans and animal models such as rodents or zebrafish^8^. Until recently, available technology in precision delivery of radiation to the heart (or subregions of the heart) in small mammalian models such as mice posed another difficulty in studying RICT in animals, although recent developments in partial heart irradiation have now been reported^9^. Additional limitations due to atypical responses in available mouse strains (for example, C57 sub-strains present with markedly distinct responses in terms of fibrosis and pleural effusions following thoracic irradiation^10^) mean that the biological mechanisms underlying RICT remain incompletely understood^11^. There is an urgent need to further understand the mechanisms that underlie RICT using appropriate models and thus develop potential cardioprotective therapeutic approaches.

The A/J mouse has been shown to develop relevant pathophysiological adverse events to radiotherapy including pneumonitis following thoracic radiation therapy^12^. We previously reported development of inflammatory oesophagitis in the A/J mouse approximately one week after thoracic irradiation^13^. The development of inflammation-induced events following irradiation suggest that the A/J mouse is a useful model for the evaluation of other early and late toxicities from radiotherapy. The aim of the present study was to investigate potential mechanisms that underlie normal cardiac tissue responses to radiotherapy and development of RICT using the A/J mouse as a model.

## Methods and Materials

All animal work was performed under project license 30/3395 issued by the UK Home Office. Female A/J mice (aged 6 – 8 weeks) were purchased from Envigo (UK) and housed in ventilated cages. Housing was maintained 22°C and 55% humidity with 12 h dark and light cycles.

### Electrocardiogram

Mice were anaesthetised using isoflurane (2-4%) and body temperature was maintained at 37°C and respiration rate was monitored and maintained at 40–60 breaths/mins during the course of data collection. Needle electrodes were inserted subcutaneously under the skin of the upper abdomen of the mice to collect lead I surface electrocardiogram (ECG) traces. Traces were acquired with PowerLab (ADInstruments, UK) under a controlled isoflurane dose (2-4%) and stable breathing for a duration of 5-10 minutes. ECG readings were acquired 22 and 26 weeks after thorax or heart irradiation.

ECG traces were analysed for R-R intervals and waveform segmentation using LabChart 8 (v8.1.13). An ROI consisting of 100-300 consecutive heartbeats were selected on each ECG trace. Traces with an isoelectric noise < 0.1 mV and < 8 mV activity, and R-R intervals < 0.2 seconds and a form factor < 125 were excluded from the analysis. An ECG analysis module (LabChart 8, ADInstruments) was applied with mouse ECG trace settings (4 beats averaging; pre-P baseline of 10ms, maximum PR of 50 ms and maximum RT of 40 ms) for segmentation of ECG waveforms to identify P-wave, QRS complex and T-waves. Heart rate variability (HRV) was assessed as the root mean square of successive differences (RMSSD) between heartbeats. R-R interval values were obtained from analyses carried out with the ECG analysis module with settings specified as above, and RMSSD was calculated using the formula below.

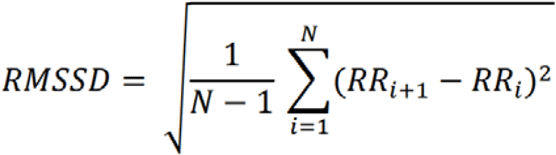

### Irradiation of the heart

Mice were anaesthetised (2-4%) and placed on a mouse bed for 360° image acquisition with computer tomography (CT) using an image-contouring step with a CT2MD system. Radiation beam size and position were calculated using the Small Animal RadioTherapy (SmART) Plan treatment system and an X-ray was delivered with an arc beam on the X-Rad225Cx small animal irradiator (Precision Xray, North Branford CT) at 10-400 cGy/min (225kV, 13 mA). Dose planning and irradiation were performed using machines and facilities at Suzhou University (Suzhou, China) by researchers at SuZhou University and Crown Bioscience (Suzhou, China).

### Histology and immunohistochemistry (IHC)

Heart and lungs were isolated from the mice and formalin-fixed and paraffin-embedded. Central transverse sections of the heart were cut into 4 μm sections and stained with haematoxylin and eosin (H&E) and Masson’s Trichrome staining (#HT15, Sigma-Aldrich, UK) for collagen detection. IHC was performed using γH2AX Ser139 (Cell Signaling Technologies, 9718) and phospho-p53 ser15 (Cell Signaling Techologies, 9284) antibodies and the Dako EnVision+ G2 Doublestain System (K4003 and K3468, Agilent) per the manufacturer’s instructions. Histology sections were scanned with the Aperio CS scanner and QuPath (v0.4.3) was used for image analysis.

### AI-based digital pathology for collagen detection

Collagen thickening around the perivasculature was quantified by the area of collagen tissue per μm of blood vessels. The miniNet AI algorithm (HALO Image Analysis Platform Software, Indica Labs) was used to identify collagen tissue, myocardial tissue and glass on Masson’s Trichrome-stained images of cross-sections of the heart. The AI was trained by at least 15 manually annotated regions of collagen, myocardium and glass. The calculated total area of collagen was used for quantification.

### Second-harmonic imaging microscopy

4 μm longitudinal heart sections were stained with haematoxylin and eosin using standard methods and mounted with DPX. One whole-tissue image and three to five ROIs were imaged (including the pericardium in the field) per sample on the Zeiss 880 NLO confocal microscope using the Mai Tai Ti: Sapphire oscillator (Spectra-Physics, California, USA). SHIM Images were captured with an excitation wavelength of 890 nm, a pulse width of less than 100 fs and an emission filter centred at 445 nm and 10x objective. Microscopy images acquired on the Zeiss 880 microscope were deconvoluted on the ImageJ software. Pericardial regions were manually labelled on individual images and an algorithm-based image analysis tool on ct-FIRE software (https://loci.wisc.edu/software/ctfire) was used to analyse collagen fibre length, width, straightness and angles.

### Flow cytometry

Mice were perfused with ice-cold Hanks Balanced Salt Solution (HBSS). Hearts were collected and atria were removed. Single cells were obtained by mincing and digesting the tissues with collagenase/DNase solution followed by red blood cell removal. Cardiac cells were blocked with anti-mouse CD16/32 antibody (BioLegend, 101302) and stained with live-dead stain (Invitrogen, L23101) and anti-mouse CD45 (BioLegend, 103116), F4/80 (BioLegend, 123177), CD11b (BioLegend, 101222), CD80 (BioLegend, 104707) and CD206 (BioLegend141707) antibodies. Flow cytometry was performed on the CytoFLEX Flow Cytometer (Beckman Coulter, Indianapolis US) and analysis was carried out using FlowJo (v10.6.1, BD Bioscience).

### Drug treatments

Where relevant, urethane (SigmaAldrich) was administered in PBS at 5% w/v as a single dose intraperitoneally. DMXAA (SelleckChem) was administered in 5% DMSO, 5% NaHCO3 in water at 10 mg/kg as a single dose intraperitoneally.

### Statistical analysis

Data were expressed as mean ± standard error of the mean (SEM) and analysed using the GraphPad Prism software (version 9.0, GraphPad). Student’s t-test was used for comparison of two groups, and one-way analysis of variance (ANOVA) with a Bonferroni correction was used on comparisons of more than two groups unless otherwise specified. Statistical significance was defined as ^ns^p > 0.05, *p < 0.05, **p < 0.01, ***p < 0.001, ****p < 0.0001.

## Results

### A/J mice develop cardiac toxicity following whole heart irradiation

To first evaluate the A/J mouse model for the studies of radiation-induced cardiac toxicity, A/J mice were treated with CT-guided whole-heart RT at a single dose of 13 Gy or 15 Gy (mock RT for control mice). To evaluate late cardiotoxic effects, ECG traces were collected 6 months after RT and mice euthanised for analysis of cardiac tissue (**Figure 1a**). Sudden death, pleural effusion, ventricular dilation, arrhythmias, and pericardial fibrosis were observed in the A/J mice that were given a dose of RT to the heart. Specifically, pleural effusion was observed in 33.3% of mice treated with 13 Gy RT and 50% of mice treated with 15 Gy RT upon necropsy at the experiment endpoint (**Table 1**). Notably, a case of sudden death was observed 17 weeks after 15 Gy RT (1 out 6 mice; 16.7% of group), while all mice survived until the experimental endpoint in the 13 Gy RT and mock RT groups.

**Table 1.** Clinical and pathophysiological phenotypes of late radiation-induced cardiac effects in A/J mice.

|  | **No RT** | **13 Gy (heart)** | **15 Gy (heart)** |
| --- | --- | --- | --- |
| **Sudden death** | 0% (0/3) | 0% (0/3) | 16.7% (1/6) |
| **Pleural effusion** | 0% (0/3) | 33.3% (1/3) | 50% (3/6) |
| **Ventricular dilation** | 0% (0/3) | 0% (0/3) | 50% (3/6) |
| **Arrhythmia** | 0% (0/3) | 0% (0/6) | 16.7% (1/6) |
| **Pericardial fibrosis\*** | 0% (0/3) | 100% (3/3) | 100% (6/6) |

\* > 50% of pericardium with $\geq 5\mu\text{m}$ collagen at the pericardium with Masson's Trichrome stain

**Figure 1.**
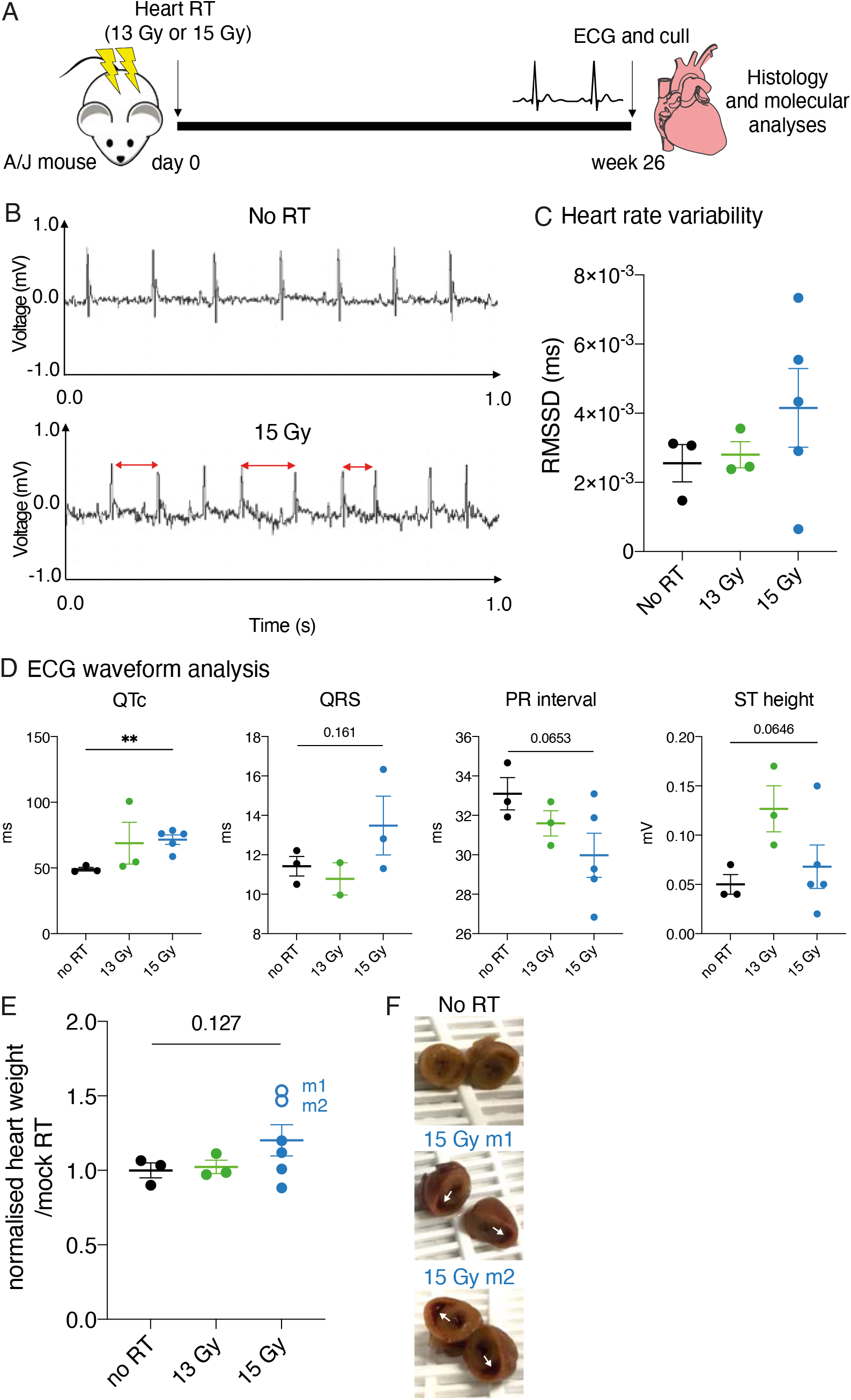
Radiation treated A/J mice demonstrate abnormal cardiac conductivity and hypertrophic cardiomyopathy. **(A)** Schematic representation of the experimental design for cardiac irradiation of A/J mice. Three to five mice were given mock, 13 Gy or 15 Gy targeted RT to the heart, six months after which ECG were acquired and mice were euthanised for analysis of heart tissue. **(B)** Representative 1.0-second-long ECG traces acquired from A/J mice 26 weeks after mock and 15 Gy RT. One of six mice in the 15 Gy cardiac irradiated and two of six mice in the 13 Gy thorax irradiated mice developed arrhythmia six months after RT. **(C)** Heart rate variability was measured by root mean square of successive RR interval differences (RMSSD) of mock, 13 Gy or 15 Gy heart irradiated mice. Each data point represents the mean RMSSD calculated from analysis of ECG traces of more than 100 successive heartbeats. **(D)** Mean QTc, QRS, and PR lengths and ST height calculated from an ECG analysis of more than 100 successive heartbeats acquired from mock, 13 Gy or 15 Gy RT mice. **(E)** Heart weights of mock, 13 Gy or 15 Gy heart RT mice normalised to whole-body weights. Data points indicating mice with the highest heart weights are shown in empty circles. **(F)** Pictures of transverse cross-sections of formalin-fixed hearts of mock RT mice (top panel) and 15 Gy RT mice with the highest body weights (bottom panel) shown. Enlarged right ventricles are indicated in white arrows. Results are mean ± SEM, analysed by a one-way ANOVA test. ^ns^p > 0.05, *p < 0.05, **p < 0.01, ***p < 0.001, ****p < 0.0001.

Assessment of electrical conduction of the irradiated hearts demonstrated an increase in frequency of arrythmias in the irradiated mice. While all mock RT mice had normal sinus rhythm, a case of arrythmia (16.7% of mice) was recorded in the 15 Gy RT group six months after treatment (**Figure 1b**). Upon analysis of the variability of heart rates with root mean square of successive differences (RMSSD), a numerical increase in mean heart rate variability was observed (**Figure 1c**), suggesting that a dose of 15 Gy to the heart can induce abnormalities to the electrical conduction of the heart. However, this was not statistically significant due to variability between individual mice readings. Furthermore, a detailed analysis of ECG waveforms revealed increases in QTc (p < 0.01) and QRS (p = 0.161) lengths and decreases in PR interval (p = 0.0653) and ST height (p = 0.0646) (**Figure 1d**). The prolonged QTc and QRS indicate a delay in ventricular repolarisation and is associated with an increased risk of arrhythmias and sudden cardiac death, with shortened PR interval indicating conduction delay between the right and left atria. Together, these indicate that 15 Gy irradiation caused marked alterations in electrical conduction in our mouse model, consistent with that observed in large clinical cohort studies^14^.

### Cardiac fibrosis following radiotherapy in A/J mice

Analysis of heart organ weights indicated heavier organ weights after irradiation: 50% (3 of 6) mice treated with 15 Gy had heart weights in the range of 1.2-1.5-fold change from mock RT baseline (**Figure 1e**). The same mice had observable enlargement in ventricular volumes upon cross-sectioning of the heart (**Figure 1f**). Therefore, we further investigated late cardiac effects in the heart following RT for pathological changes consistent with fibrosis. Hearts of the mice treated with 13 Gy or 15 Gy were analysed for collagen using Mason’s Trichrome staining. A thickening of collagen around pericardial and perivascular structures of the irradiated hearts were observed on Masson’s Trichrome images which were not observed in mock RT controls (**Figure 2a**). Assessment of the area of thickened pericardium confirmed the significant increase in collagen thickening in hearts of the mice treated with 13 Gy and 15 Gy compared to the mock RT control (p < 0.0001) (**Figure 2b**). Furthermore, the impact of radiation on perivascular fibrosis were dose-dependent, with mice treated with 15 Gy RT demonstrating increased perivascular fibrosis compared to 13 Gy RT (p < 0.05) (**Figure 2b**).

**Figure 2.**
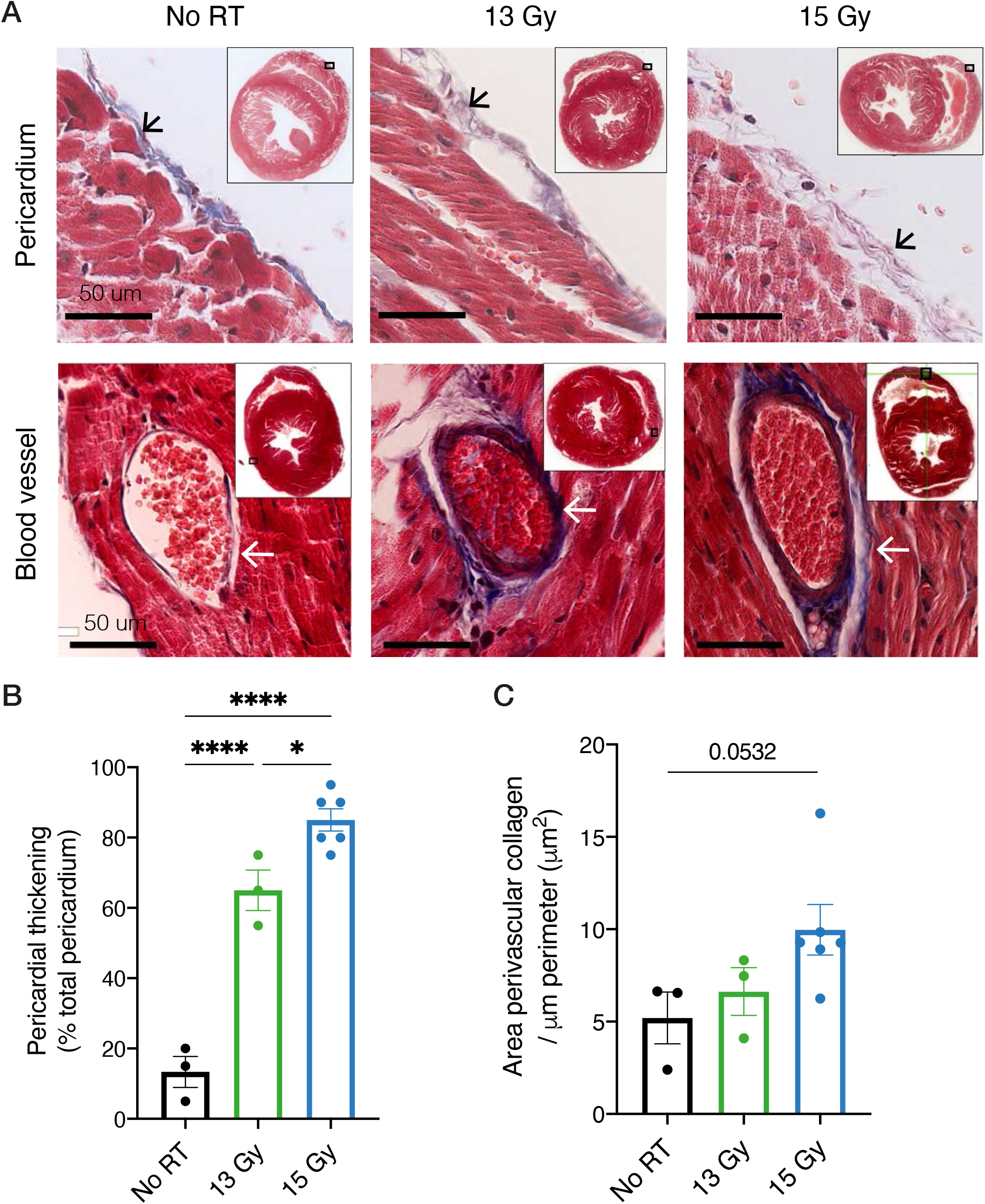
Pericardial and perivascular fibrosis post-radiation treatment of A/J mice. **(A)** Masson’s Trichrome stains of transverse cross-sections of mock, 13 Gy and 15 Gy irradiated hearts. Top: representative images of collagen in the pericardium of each treatment group. Pericardium indicated in black arrows. Bottom: representative images of collagen in the perivasculature of each treatment group. Perivasculature indicated in white arrows. Scale bars: 50 μm. **(B)** Manual pathological quantification of percent area the thickened pericardium (collagen layer ≥ 5μm) occupies of the total ventricular pericardium. **(C)** Area of perivascular collagen per perimeter of the blood vessel using a bespoke machine-learning algorithm on HALO software. Results are mean ± SEM, analysed by a one-way ANOVA test. ^ns^p > 0.05, *p < 0.05, **p < 0.01, ***p < 0.001, ****p < 0.0001.

Perivascular collagen was quantified using AI-guided tissue segmentation which identified an increase in perivascular collagen with increasing doses of radiation (p = 0.0532) (**Figure 2c**). This is consistent with recent findings of marked perivascular fibrosis, as opposed to generalised fibrosis, following irradiation, although at notably higher doses of irradiation (partial heart irradiation of 40 Gy)^9^. At the lower dose of irradiation used in this study it appears that perivascular collagen deposition was less marked, supporting the need to protect sensitive cardiac tissues from radiation exposure.

To investigate collagen organisation, second-harmonic imaging microscopy (SHIM) was performed on sectioned hearts. SHIM on mock and 13 Gy RT treated hearts demonstrated high signal intensity in atrial structures in both mock RT and 13 Gy RT groups (indicated in the white dotted circle in **Supplementary Figure 1a**). This is expected, as atria largely consist of supportive structures including collagen. SHIM also revealed differences in photon intensity between mock and 13 Gy RT groups, notably at the pericardium, and potentially across the myocardial region (pericardium indicated in white arrows in **Supplementary Figure 1a**). Analysis of individual collagen fibres was performed using the ct-FIRE software. Results did not reveal any significant differences in length and angles of the collagen fibres. However, showed a trend toward increasing disorganisation of collagen (reflected in reduced straightness and increased width) after RT (**Supplementary Figure 1b**). While similar observations have been made in normal tissue following bladder and rectal irradiation^15^ this is the first such reported analysis in mouse cardiac tissue following irradiation.

### Persistent DNA damage response in cardiac myocytes

To investigate the underlying mechanisms that could lead to late cardiac toxicities from radiation, molecular and histological analyses were performed on cardiac tissues collected at 0 hours, 2 hours, 24 hours, and 7 days after treatment with 5 Gy. IHC analysis for markers for DNA damage response (DDR) including γH2AX and phospho-p53 surprisingly revealed that DDR in the cardiomyocytes persisted at least 7 days after irradiation (**Figure 3a**). γH2AX positive nuclei increased significantly 2 hours after RT (p < 0.0001) and persisted at a similar level 7 days after exposure to 5 Gy RT (**Figure 3b**). Similarly, p-p53 positive nuclei also were significantly increased 2 hours after RT (p < 0.05) without returning to baseline levels even after 7 days (**Figure 3b**). Interestingly, studies have shown that adult cardiomyocytes have a limited capacity to re-enter the cell cycle^16,17^, however, this does not account for the persistent DNA damage response markers we observed in this cell type. Importantly, in the context of heart failure, persistent unrepaired DNA breaks have been reported using an *in vivo* model with subsequent inflammatory gene changes consistent with the induction of senescence and the senescence-associated secretory phenotype (SASP), which may account for the changes we observed in the A/J model following irradiation^18^.

**Figure 3.**
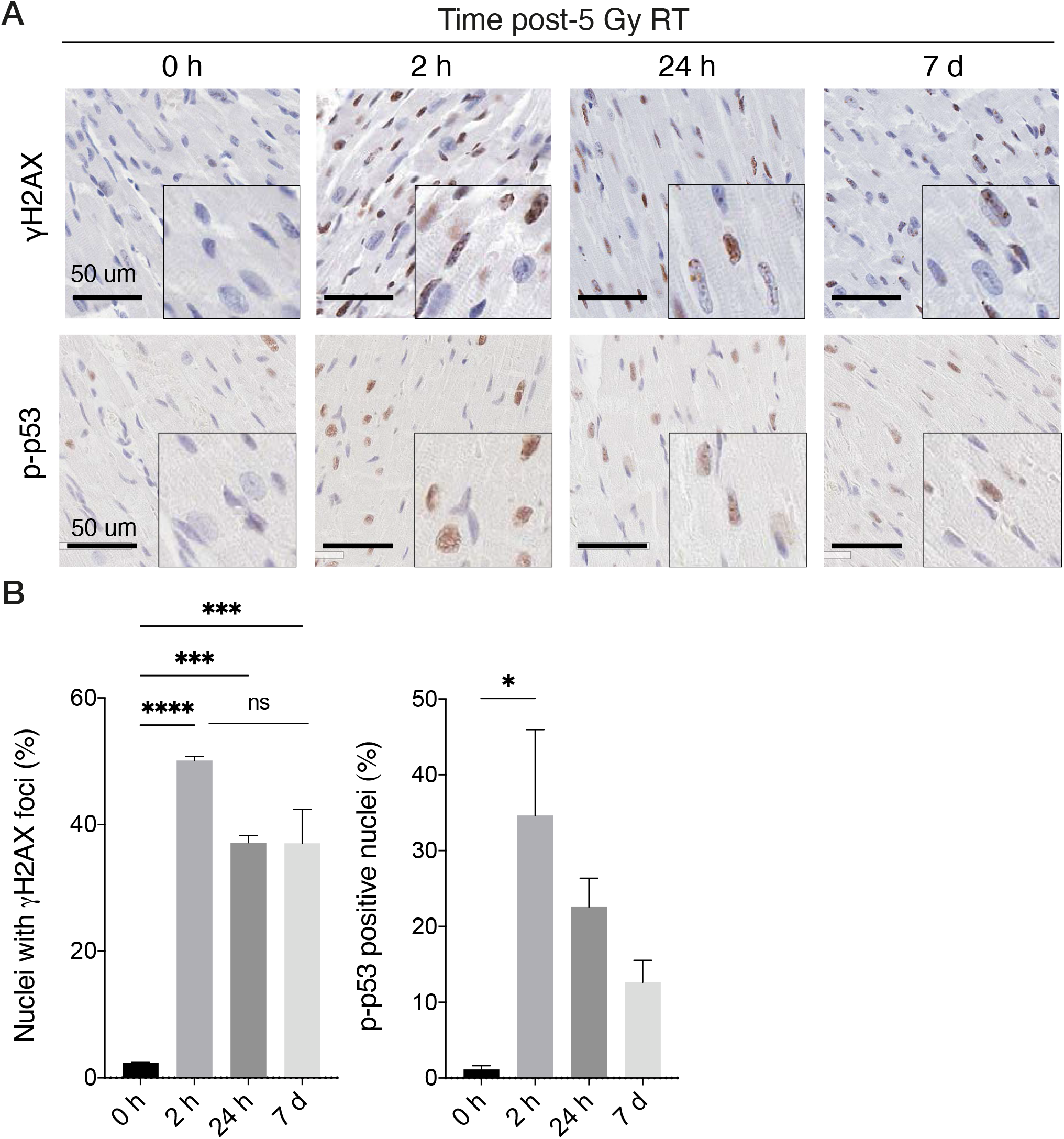
Persistent DNA-damage response in the heart is a potential contributor to radiation-induced cardiac fibrosis. **(A)** Immunohistochemistry for γH2AX and phospho-p53 on hearts was collected at 0 hours, 2 hours, 24 hours and 7 days after 5 Gy radiation treatment (n = 3 per group). Scale bars: 50 μm. **(B)** Quantification of nuclei positive for γH2AX foci and cells positive for phospho-p53 on IHC images. Each data point represents a mean value from an analysis of five ROIs per section (with a minimum of 50 nuclei per ROI). γH2AX foci positive nuclei were manually counted per field, and p-p53 positive nuclei were quantified using a nuclear detection algorithm on the Aperio ImageScope software. Results are mean ± SEM, analysed by a one-way ANOVA test. ^ns^p > 0.05, *p < 0.05, **p < 0.01, ***p < 0.001, ****p < 0.0001.

### The innate immune response to DNA damage in cardiac toxicity

Persistent genotoxic stress and the accumulation of DNA damage can result in micronuclei formation and cytosolic DNA,^19–22^which can subsequently lead to the activation of type I IFN and other innate immune pathways. Therefore, we hypothesised that the persistent DNA damage repair response and activation of innate immune pathways could contribute to fibrosis and cardiac toxicity through DNA sensing pathways such as the cGAS-STING pathway. cGAS-STING pathway activation has recently been reported as a result of cardiac irradiation^23^ and cGAS-STING signalling is implicated in the pathophysiology of many cardiovascular diseases including heart failure^24^.

To investigate if IFN stimulation alone impacts cardiac fibrosis, we treated A/J mice with a single dose of STING agonist, DMXAA (Vadimezan), intraperitoneally (i.p.) with or without 13 Gy RT. These animals were also administered urethane – we previously confirmed that urethane alone does not impact cardiac fibrosis (**Supplementary Figure 2**). Mice were euthanised and assessed for cardiac fibrosis 26 weeks after treatment (**Figure 4a**). DMXAA treatment alone resulted in mild increased pericardial fibrosis (p < 0.05). When DMXAA was combined with RT there was a marked increase in cardiac fibrosis (p < 0.0001) although this was not significantly increased in comparison to RT alone (**Figure 4b, c**). Systemic STING activation therefore did not appear to significantly increase RICT.

**Figure 4.**
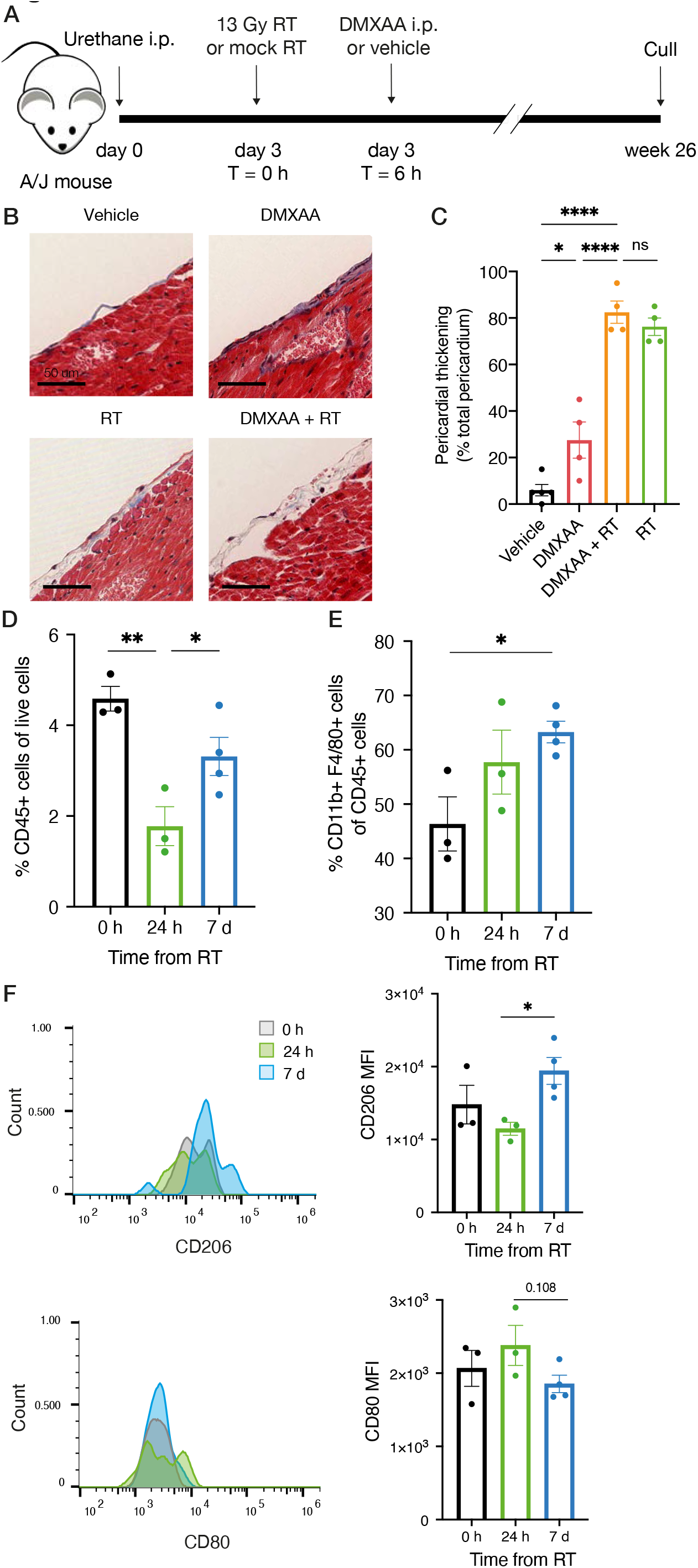
Pericardial fibrosis and innate immune response to radiation therapy. **(A)** Schematic representation of the experimental schedule for DMXAA and radiation treatment of A/J mice. **(B)** Pericardial thickening quantified from Masson’s Trichrome-stained hearts collected 24 weeks after DMXAA +/-RT and vehicle/mock RT treatments. Results are mean ± SEM, analysed by a one-way ANOVA test. ^ns^p > 0.05, *p < 0.05, **p < 0.01, ***p < 0.001, ****p < 0.0001. **(D)** Flow cytometry analysis of macrophage populations 0 hours, 24 hours, and 7 days after 13 Gy thoracic RT. Percent CD45+ cells of live cells 0 hours, 24 hours and 7 days post-RT. **(E)** Percent CD11b+ F4/80+ macrophages of CD45+ cells 0 hours, 24 hours and 7 days post-RT. **(F)** Mean fluorescence intensity (MFI) of CD80 and CD206 in macrophage populations selected with CD11b+ F4/80+ gating on flow cytometry. Top: histogram of CD206+ M2-like populations in cardiac tissue 0 hours, 24 hours and 7 days after RT, and quantification of MFI. Bottom: histogram of CD80+ M1-like populations in cardiac tissue 0 hours, 24 hours and 7 days after RT, and quantification of MFI. Results are mean ± SEM, analysed by a one-way ANOVA test. ^ns^p > 0.05, *p < 0.05, **p < 0.01, ***p < 0.001, ****p < 0.0001.

### Immune infiltration in cardiac tissues following irradiation

To investigate if the innate immune system is affected by RT treatment, flow cytometric analyses of cardiac tissue was performed at 0 hours, 24 hours, and 7 days after 13 Gy RT. Data revealed that there was an overall decrease in CD45^+^ lymphocytes 24 hours after irradiation and recovery of the CD45^+^ population to baseline (0 hours) 7 days after RT (p < 0.05) (**Figure 4d**). The F4/80^+^ CD11b^+^ myeloid population makes up 46.4% of the CD45^+^ population at baseline and, in contrast, this population increases significantly 7 days after irradiation (p < 0.05) (**Figure 4e**). Furthermore, analysis of the F4/80^+^ CD11b^+^ myeloid population with CD206 mean fluorescence intensity (MFI) and CD80 MFI revealed an increase in CD206 MFI between 24 hours and 7 days after RT (p < 0.05), and a trend toward a decrease in CD80 MFI between 24 hours and 7 days after RT (**Figure 4f**). These data suggest that there is a shift away from an M1-like phenotype (identified by CD80) and toward an M2-like (identified by CD206) phenotype in the macrophage populations in the irradiated heart.

## Discussion

Here we have characterised the A/J mouse model to study the early and late molecular and pathological effects of RICT can be studied, either alone or in combination with other systemic therapies. In keeping with previous reports, late RICT were characterised as pericardial and perivascular fibrosis and electrophysiological abnormalities, as well as changes in ventricular structure. We also identified several events occurring hours to days after RT that could be contributing to RICT in the A/J mice. These include persistent DNA-damage response as shown by sustained γH2AX foci and p-p53 expression and early macrophage polarisation toward a CD206+ M2-like phenotype. We hypothesise that some of the immunological changes that we see happening (e.g., M2 repolarisation) could be contributing to the accumulation of collagen around damaged tissue. However further investigation is needed to mechanistically identify the relationship between innate immune responses and collagen deposition.

Radiation-induced abnormalities in electrical conduction are currently an area of interest, and although epidemiological studies have suggested this correlation exists^25–27^, thus far electrical abnormalities following irradiation have not been described in mouse models. Therefore, our study adds to the existing knowledge on the impact of cardiac RT on the electrical output and potential development of cardiac arrhythmias as we identified RT as associated with increased variability in the heart beats and waveform changes. Abnormality in the electrical conduction could have led to the case of sudden death we observed in the 15 Gy heart RT treated animals, as we also see an overall increase in abnormal conduction (greater heart rate variability and QTc/P-wave changes) in the 15 Gy RT condition.

Interestingly, a recently reported study^23^ demonstrated that treatment with a STING antagonist could suppress IFN responses and subsequent cardiac toxicity induced by DNA-damaging agents (including doxorubicin and RT) and also that RICT was reduced in cGAS and STING KO mouse models. This study observed persistent DNA damage signalling in endothelial cells, not cardiac myocytes, based on RNAseq analysis. In contrast, using IHC analysis of H2AX and p-p53, we observed persistent DDR in cardiac myocytes. A limitation is that a myocardial cell maker (e.g. ACTN2) was not included in our analysis, and that distinct modalities (RNAseq compared to IHC analysis) were employed. However, our findings are consistent with those reported in the context of a model of heart failure, where persistent unrepaired DNA damage in cardiac myocytes was associated with inflammatory gene changes and potentially activation of senescence and SASP^18^. Further study of SASP as a potential mechanism contributing to inflammation and fibrosis following cardiac irradiation is warranted.

The location in which fibrosis was most pronounced months after RT – pericardium and perivasculature – suggests the possibility that structures that are closer to blood vessels where immune cells can infiltrate are more vulnerable to fibrotic events than the myocardium that are further away from these structures, for instance. The timing in which cardiac fibrosis is observed in the A/J hearts - approximately six months after RT – suggests that the acute events (DDR and immune events) that we observed in our studies could be acting as the initial onset for later, more gradual processes to occur in the cardiac tissues over a long period, potentially via the induction of SASP. This model could also potentially be used to study physiological phenotypes and valvular alterations following RT to the heart.

In summary, the A/J mouse responds in a physiologically relevant manner to cardiac irradiation and can be used to study potential mechanisms driving RICT. We identified persistent unrepaired DNA damage in cardiac myocytes and a potential role of STING-driven interferon, recently also reported by others^23^, in driving inflammation and fibrosis. Further studies are required to further delineate the relationship between persistent DNA damage, SASP, macrophage repolarisation and collagen deposition to identify potential therapeutic strategies, with our findings an important contribution to deeper understanding of the causative factors of RICT in patients treated with radiotherapy.

## Supplementary Figures

**Supplementary Figure 1.**
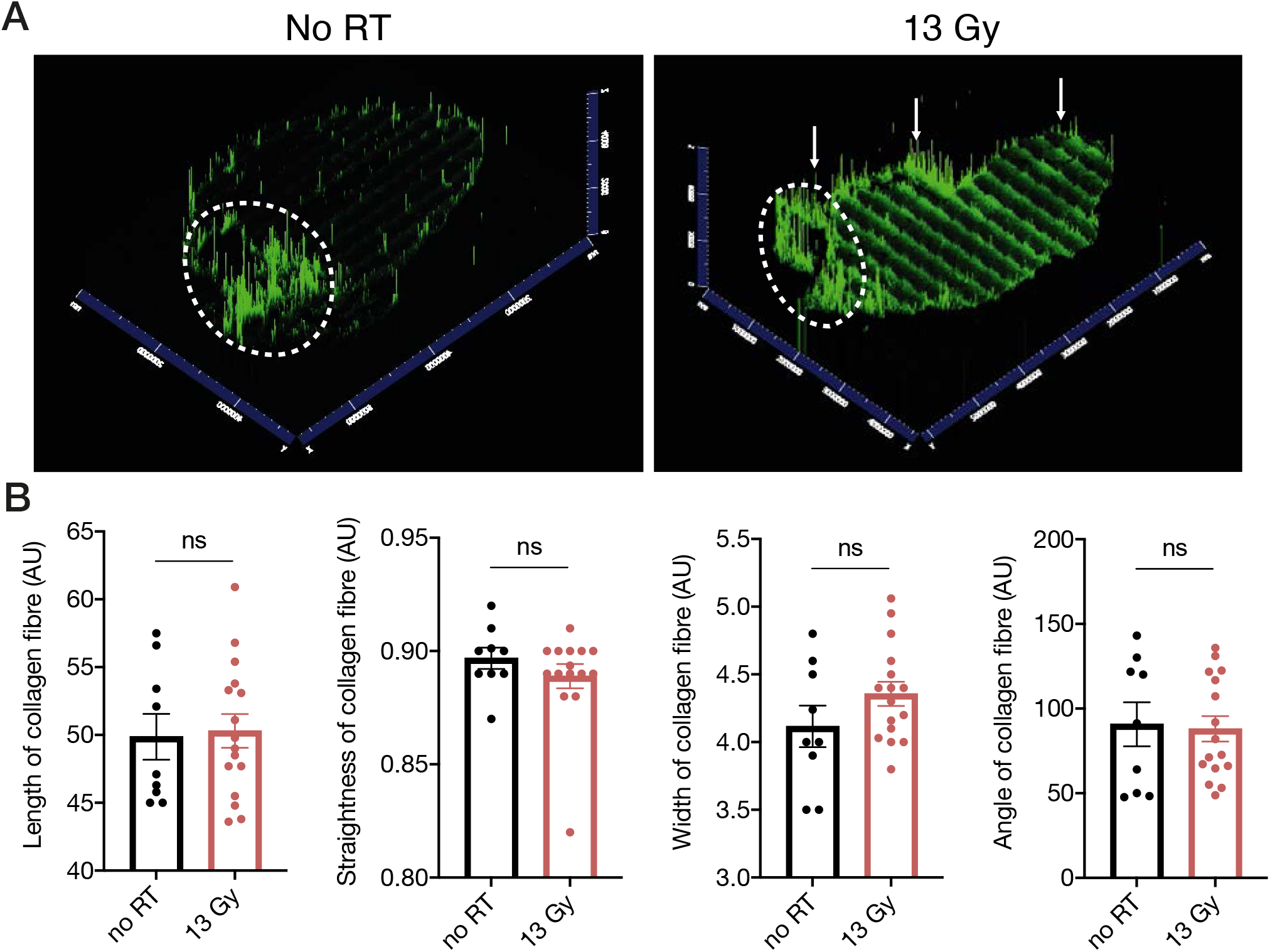
SHIM analysis does not indicate change in length or width of collagen the pericardium after radiation treatment. **(A)** Representative images of longitudinal cross-sections of mock irradiated and 13 Gy irradiated hearts imaged by second-harmonic microscopic imaging (SHMI). Atria are indicated in dotted white circles, and pericardium is indicated in white arrows on the 13 Gy irradiated heart section. **(B)** Quantification of length, width, and straightness of collagen fibres of mock and RT treated heart sections. SHMI of three whole-heart sections were collected per group. Images were deconvoluted and three to five ROIs including the pericardium were selected per section and analysed using the ct-FIRE software. Each data point represents a value generated per ROI. Error bars indicate SEM. Student t-test. *p<0.0001.

**Supplementary Figure 2.**
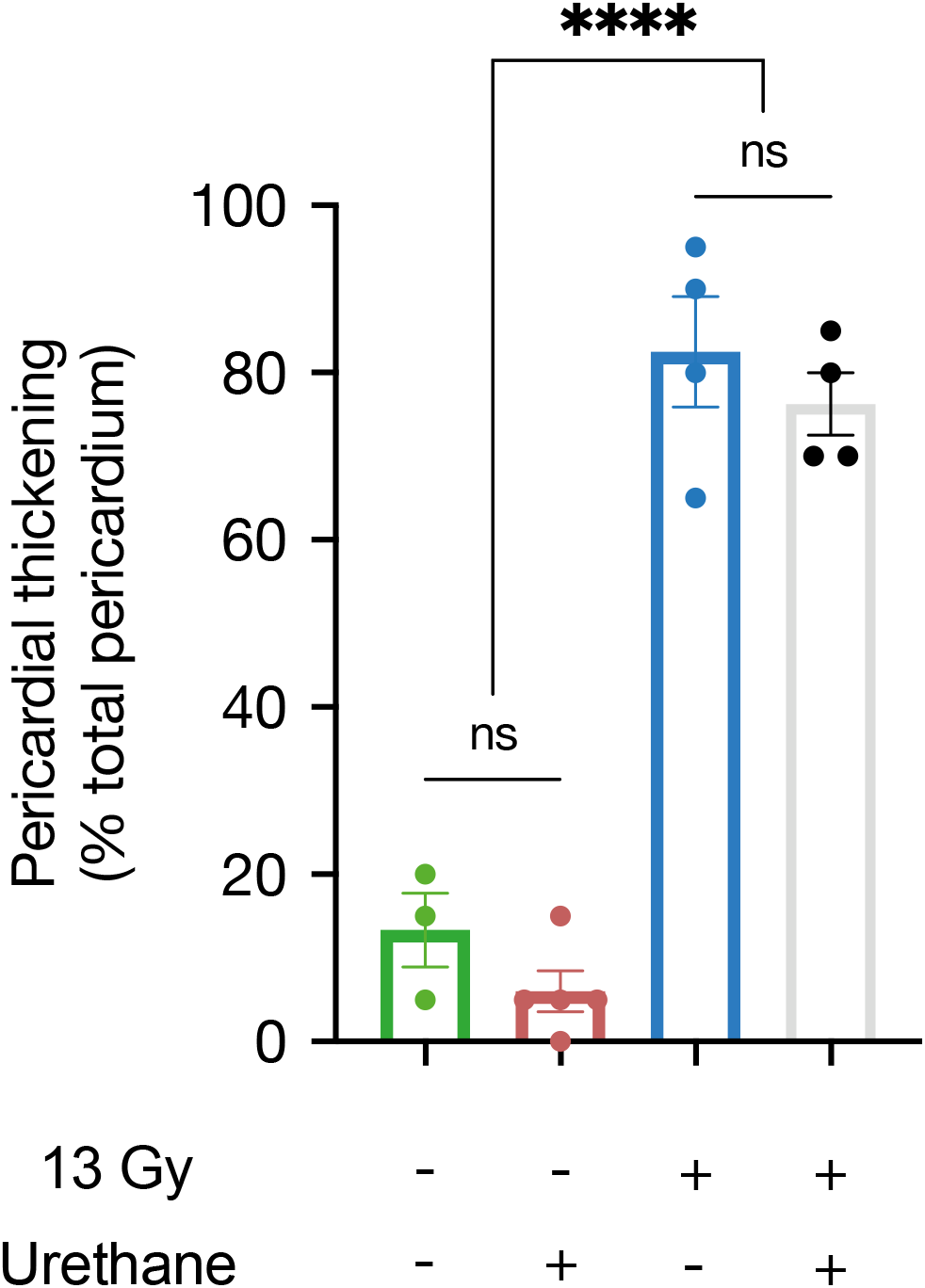
Urethane does not increase cardiac fibrosis. Pericardial thickening quantified from Masson’s Trichrome-stained hearts collected 24 weeks after RT/mock RT +/-urethane pre-treatment. Results are mean ± SEM, analysed by a one-way ANOVA test. ^ns^p > 0.05, *p < 0.05, **p < 0.01, ***p < 0.001, ****p < 0.0001.

